# Causal reasoning is domain-specific in the human brain

**DOI:** 10.64898/2026.10.27.762213

**Authors:** R.T. Pramod, Jessica Chomik-Morales, Laura Schulz, Nancy Kanwisher

## Abstract

Causal reasoning enables humans to explain the past, predict the future, and intervene on the present. Despite extensive interest in causal inference by philosophers, cognitive scientists, and developmental psychologists, few studies have investigated causal reasoning in the human brain. Here we use fMRI to ask whether a unified brain system exists that implements causal inference across content domains, or whether instead distinct brain systems are engaged in causal reasoning about social agents and physical objects. Across three pre-registered experiments, we find that causal reasoning about social and physical phenomena engage distinct cortical regions. The fronto-parietal Physics Network (PN), previously implicated in physical reasoning, and the temporo-parietal junction (TPJ), previously implicated in social reasoning, both showed stronger responses to causal than non-causal conditions, but only within their respective domains. Selective responses emerged in both a reading comprehension task (Experiment 1) and a sentence matching task (Experiment 2), and for preventative as well as generative causal reasoning (Experiment 3). No region selectively engaged causal reasoning or showed common patterns of response across voxels for causality in both physical and social domains in whole-brain analyses. Thus, despite its central role in human cognition, causal reasoning is implemented in separate domain-specific regions rather than a single unified brain region.

## Introduction

Causal reasoning supports prediction, explanation, exploration, and intervention both in everyday life and in scientific inquiry. Little wonder that Hume described causal reasoning as the “cement of the universe”, on which “all the operations of the mind must, in a great measure, depend…” (1740). However, while causal reasoning has been a perennial topic of investigation in philosophy^1^, computer science^2^, and developmental psychology^3^, little is known about its neural bases. Here we address a question at the heart of a long-standing debate in cognitive science^4^: Does the brain have specialized machinery for causal reasoning independent of content domain, or do we engage different brain systems when reasoning about social and physical causes?

Evidence in cognitive science has been amassed on both sides^5^. Arguments for domain-generality include the facts that causal reasoning requires representing probabilistic relationships among variables independent of content domain^2,6,7^; that causal relationships of all kinds are marked with similar syntactic structures and lexical items^8–12^, and that children use identical patterns of covariation evidence to infer causal relationships whether they are reasoning about physical events (e.g., blocks making toys go) or psychological ones (e.g., silly faces making someone giggle)^13^. Arguments for domain-specificity include the facts that the computations required for reasoning about objects and forces (physics simulations) differ from those for reasoning about agents and goals (expected utilities). Additionally, both adults and children have distinct intuitive theories of physics and psychology^14,15^ and even infants have domain-specific causal representations (e.g., expecting objects to move only when contacted by an outside force but expecting agents to move on their own and act on other agents at a distance)^16–18^.

Only a few neuroscience studies have investigated causal reasoning^19–22^, and none has determined whether common or distinct brain machinery is engaged in causal reasoning across content domains. Prior work shows that distinct cortical regions in the fronto-parietal cortices (i.e., the Physics Network, or PN) are engaged in thinking about inanimate objects and their properties, relationships and events^23–27^, whereas the temporoparietal junction (TPJ)^28–30^ is selectively engaged in thinking about other people’s thoughts, and how those thoughts cause behavior^31^. One hypothesis therefore is that causal reasoning differentially recruits only these domain-specific systems. Alternatively, causal reasoning could recruit established domain-general regions such as the fronto-parietal multiple demand (MD) system^32,33^. Finally, there may be a distinct neural system that is specifically engaged in causal reasoning across content domains. Here, we tested these three hypotheses in three pre-registered fMRI experiments.

## Experiment 1: Testing Causal Reasoning in a Reading Comprehension Task

We collected BOLD responses while 31 participants read short paragraphs and answered follow-up questions (Figure 1A). Paragraph-question pairs fell into four conditions that resulted from crossing domain (physical or social) by reasoning type (causal or non-causal). We also ran standard functional localizer scans to identify in each participant individually i) the fronto-parietal “physics network” (PN)^23^, ii) the bilateral TPJ^31^, and iii) the “Multiple demand” (MD) system, which has been shown to respond during increased cognitive demand largely independent of content^32,33^.

**Figure 1.**
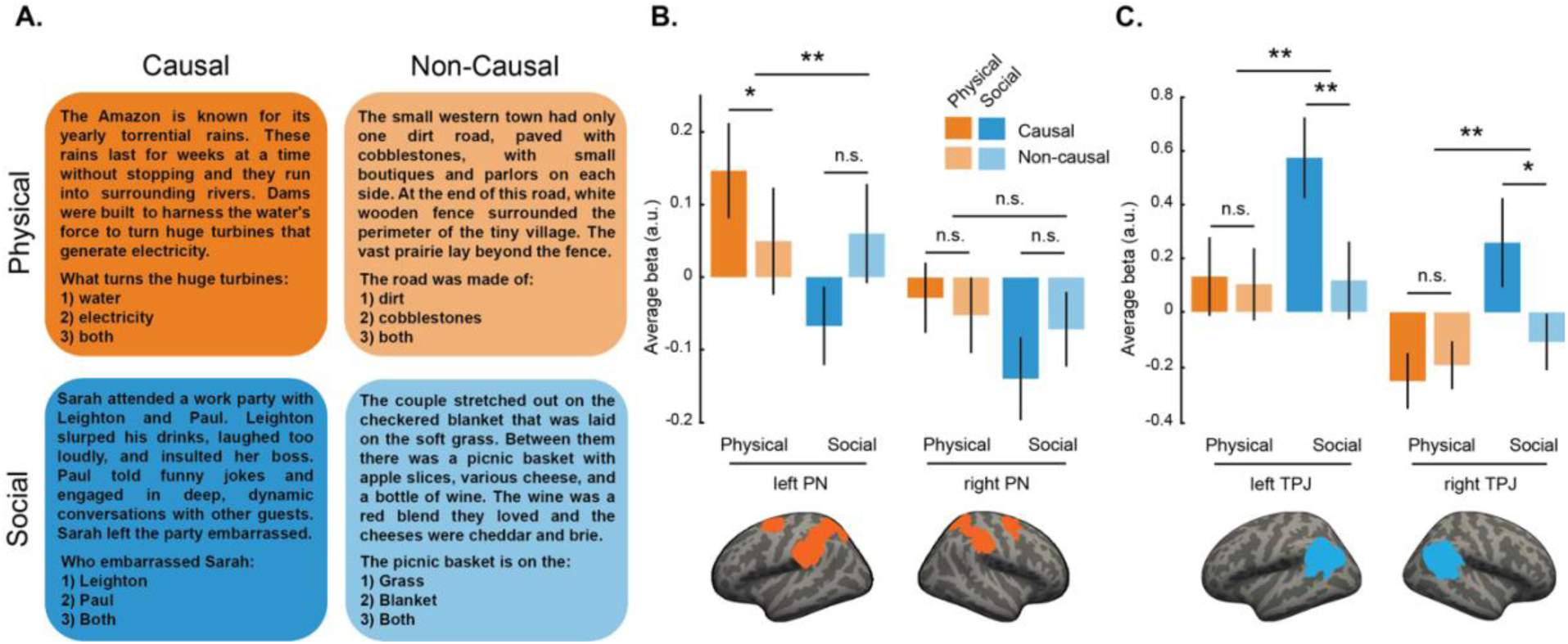
Experiment 1 (Reading Comprehension Task). (A) Example stimuli showing paragraph-question pairs from the four conditions: Physical/Social domains x Causal/Non-causal conditions. (B) Bar plot showing the average fMRI response to each of the four conditions in the left and right Physics Network (PN), N = 31. (C) Similar bar plots showing the average activation to each of the four conditions in the left and right TPJ, N = 8 (Theory of Mind localizer data was not available in other participants). The anatomical PN and TPJ parcels used for this and subsequent functional Regions of Interest (fROI) analyses are shown below the bar plots on an inflated brain surface. * is p < 0.05, ** is p < 0.005 and n.s. indicates *not significant*. Error bars indicate standard error of mean. The anatomical parcels

The domain-specific causality hypothesis predicts that physical causal reasoning should selectively engage the PN while social causal reasoning should selectively engage the TPJ, whereas the domain-general hypothesis predicts similar activations for physical and social causal reasoning, engaging either the MD system or another system. Because our stimuli are verbal (written sentences) we predicted that any of these effects may be more prominent in the left hemisphere. All hypotheses and analysis methods were pre-registered on OSF, except where indicated otherwise.

Consistent with the domain-specific hypothesis of causal reasoning, we found selective responses to the physical causal condition in the PN (Figure 1B, see Figure S7 for sub-regions), and the social causal condition in the TPJ (Figure 1C, see Figure S6 for rest of the ToM network). These findings were statistically supported by significantly higher responses in each region for i) the causal condition in the preferred domain (physical for PN, social for TPJ) than the noncausal condition in the preferred domain, ii) the causal condition in the preferred domain than the causal condition in the nonpreferred domain, and iii) a significant interaction between physical/social x causal/noncausal (see Table 1). Importantly, the domain-specific effects in both PN and bilateral TPJ could not be explained by task behavior (accuracy and reaction time; Figure S1). Additionally, stimuli were matched for word count and grade-level readability, making linguistic confounds unlikely explanations for our results.

**Table 1:**
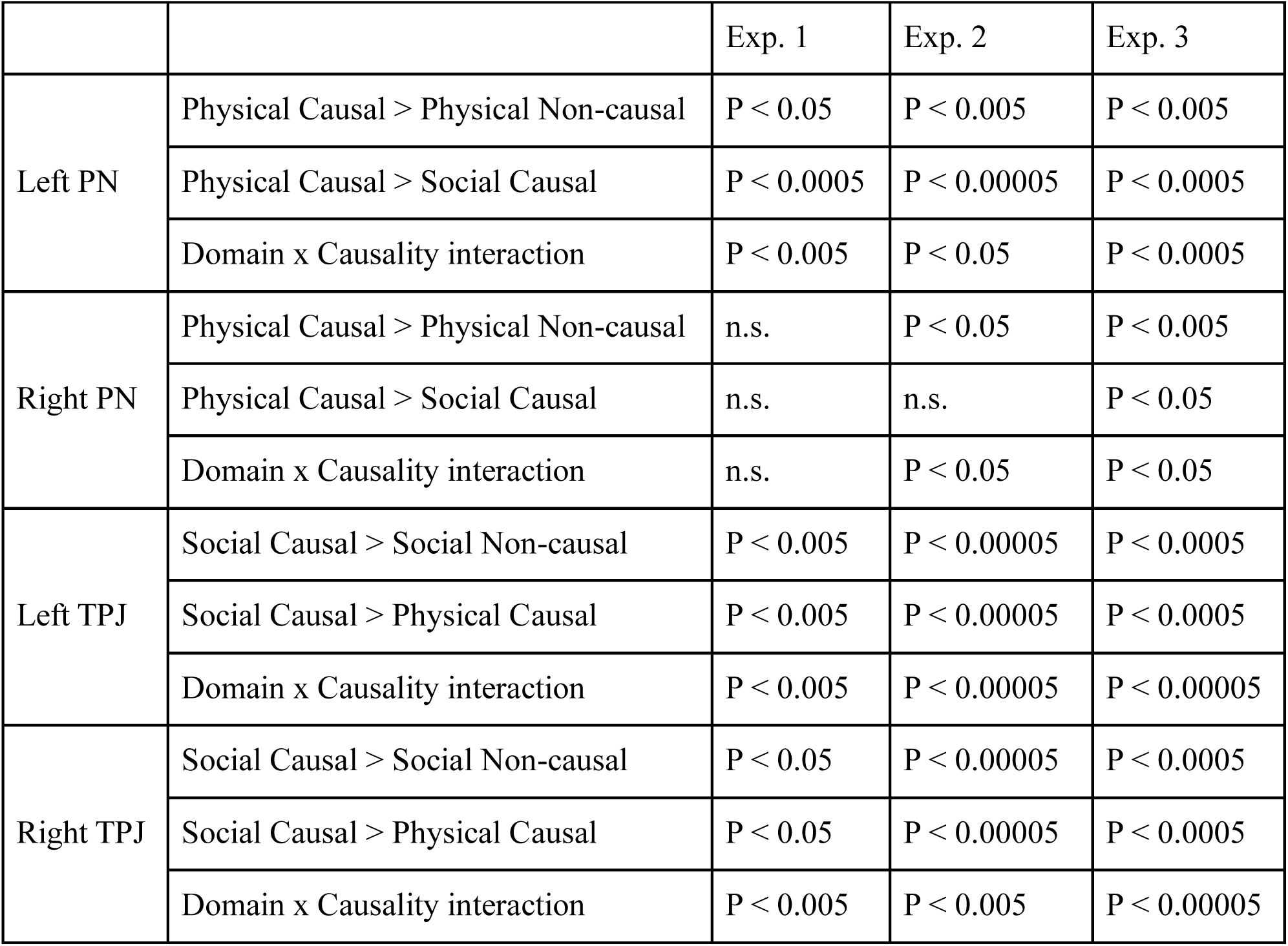
Statistical tests supported the key predictions of the domain-specific causality hypothesis in left PN and bilateral TPJ in all three experiments. All p-values are from one-sided Wilcoxon signed rank tests except the Domain x Causality interaction, which is based on repeated measures, mixed-effects ANOVA with subjects as random factors, and domain and causality as fixed factors.

|  |  | Exp. 1 | Exp. 2 | Exp. 3 |
| --- | --- | --- | --- | --- |
| Left PN | Physical Causal > Physical Non-causal | P < 0.05 | P < 0.005 | P < 0.005 |
|  | Physical Causal > Social Causal | P < 0.0005 | P < 0.00005 | P < 0.0005 |
|  | Domain x Causality interaction | P < 0.005 | P < 0.05 | P < 0.0005 |
| Right PN | Physical Causal > Physical Non-causal | n.s. | P < 0.05 | P < 0.005 |
|  | Physical Causal > Social Causal | n.s. | n.s. | P < 0.05 |
|  | Domain x Causality interaction | n.s. | P < 0.05 | P < 0.05 |
| Left TPJ | Social Causal > Social Non-causal | P < 0.005 | P < 0.00005 | P < 0.0005 |
|  | Social Causal > Physical Causal | P < 0.005 | P < 0.00005 | P < 0.0005 |
|  | Domain x Causality interaction | P < 0.005 | P < 0.00005 | P < 0.00005 |
| Right TPJ | Social Causal > Social Non-causal | P < 0.05 | P < 0.00005 | P < 0.0005 |
|  | Social Causal > Physical Causal | P < 0.05 | P < 0.00005 | P < 0.0005 |
|  | Domain x Causality interaction | P < 0.005 | P < 0.005 | P < 0.00005 |

These results also provide the first clear evidence that the PN responds selectively to physical content conveyed through language. Further, although the PN is anatomically and functionally distinct from the language network^28^, our findings indicate that linguistic input can engage domain-specific representations of physical causal relationships in the PN, mirroring prior findings that linguistic input can engage domain-specific representations of mental causes in the TPJ^31^, collectively supporting the “Exportation Hypothesis”^34^.

Confirming the domain-specificity observed in the fROI analysis, a whole-brain random-effects analysis (Figure S3) revealed significant voxels in bilateral TPJ and superior temporal sulcus (STS) for the social causal > social non-causal contrast, and significant activity predominantly in left parietal PN for the physical casual > physical non-causal contrast.

Despite the domain-specific effects reported above for the PN and TPJ, it remains possible that additional brain regions could perform domain-general causal reasoning. In exploratory analyses, we first tested for domain-general causal reasoning within the Multiple Demand (MD) network. We localized the MD network in individual participants using a spatial working memory task^33^, and measured responses to the four reading comprehension task conditions. We found a significant *opposite* trend (non-causal > causal) for both domains in the right hemisphere and only the social domain in the left hemisphere (Figure S2). Thus, we found no evidence for domain-general causal reasoning in the MD system.

We further tested for domain-general causal reasoning by the conjunction of causal > non-causal contrast in both domains (i.e, [PC > PNC] & [SC > SNC]) in a whole-brain random effects analysis. This analysis did not reveal any significant cluster of voxels (thresholded at p = 0.001, cluster-corrected) in both hemispheres (Figure 3, *top row*). A similar analysis looking for domain-specific effects found significant voxels in the parietal and temporal lobes for physical and social domains respectively, mirroring the fROI-based analyses. However, since random effects analyses pool over subject-level maps that may be spatially heterogeneous across participants, they are prone to miss genuine group-level effects. To address this issue, we performed a Group-constrained Subject Specific (GSS) analysis, which allows discovery of regions (or ‘parcels’) that are most systematically co-activated across subjects by accounting for spatial heterogeneity between individual subjects^35^. None of the domain-general parcels (i.e., regions showing significant causal > non-causal contrast in both physical and social domains) identified in one half of the data showed domain-generality in the other half of the data upon correcting for multiple comparisons.

In sum, Experiment 1 found strong evidence for domain-specific causal effects in the PN and TPJ, and no evidence for domain-general causal reasoning in the MD system or in a whole-brain univariate analysis.

The stimuli used in this experiment were designed to be naturalistic, vivid, and engaging. But their complexity makes tight control over linguistic and semantic features impossible, leaving open potential alternative accounts of our findings. We therefore designed a new experiment using a simpler stripped-down task to see if our findings would replicate when causal and noncausal reasoning were more tightly controlled.

## Experiment 2: Testing Causal Reasoning in a Sentence Matching Task

In this experiment we collected BOLD responses using fMRI while 38 participants performed a simple sentence matching task that distills causal reasoning down to its essence: matching effects to their causes. This task, which was inspired by a cause-and-effect computer game for children, had four conditions obtained by crossing two factors: domain (physical or social), and task type (matching cause and effect pairs, or matching sentences describing the same entity; see examples in Figure 2A). In each trial, the participants were instructed to find the best match for each of the four target sentences in the bottom row from four sentences in the top row of the display (see Figure 2B for an example trial). The PN, TPJ, and MD system were again localized in each participant individually using functional localizers as in Experiment 1.

**Figure 2.**
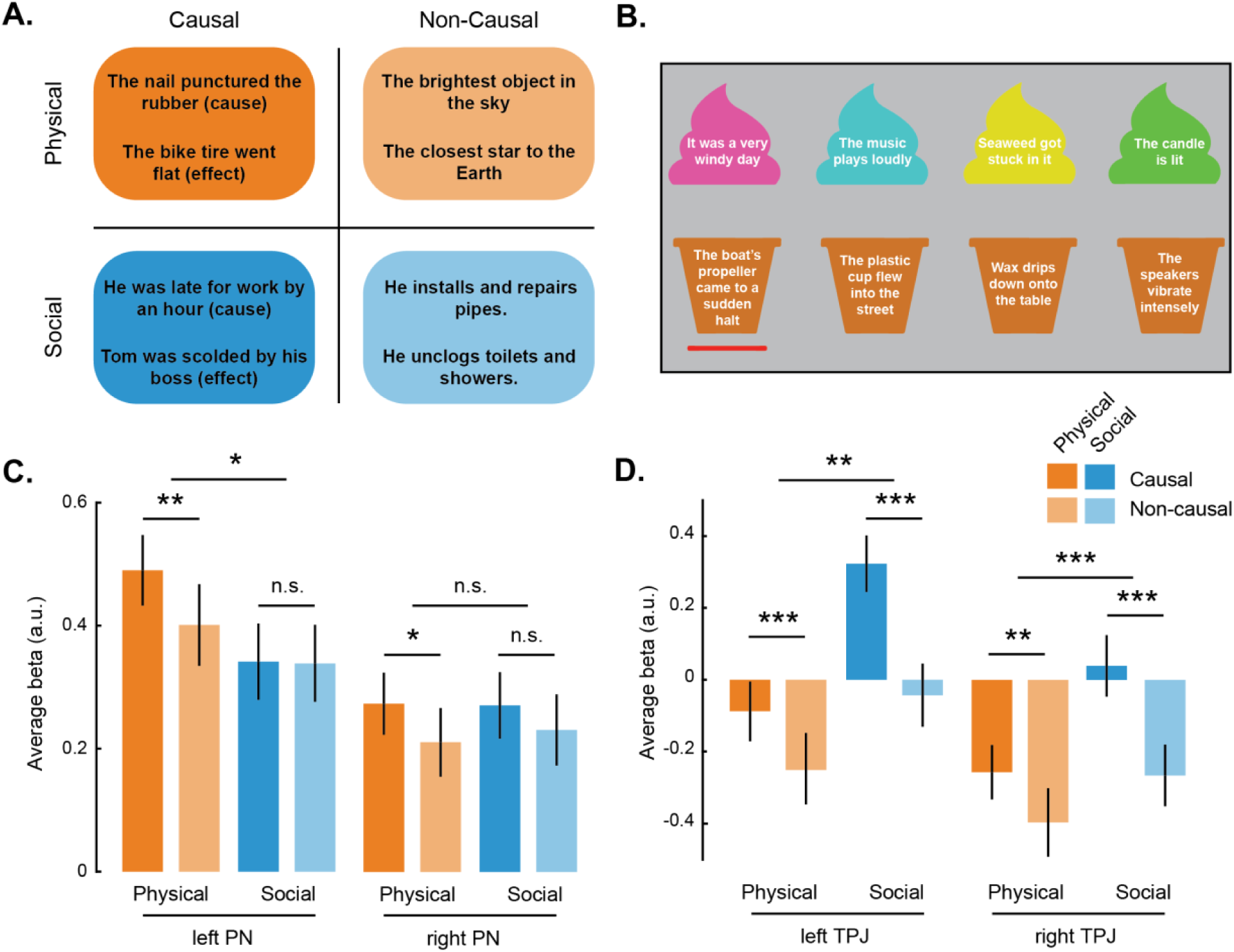
Experiment 2 (Sentence Matching Task). (A) Example stimuli showing pairs of sentences from the four conditions: Physical/Social domains x Causal/Non-causal conditions. (B) Example trial from the *Physical Causal* condition showing the causes in the top row (‘scoops’), and the effects to be matched (‘cones’) in the bottom row. The red underbar indicates the effect to which a cause must be matched; in each trial this bar moved sequentially to the right, so participants paired each effect with its cause. (C) Bar plot showing the average fMRI response to each of the four conditions in the left and right hemispheric Physics Network (PN), N = 38. (D) Similar bar plot showing the average activation to each of the four conditions in the left and right hemispheric Tempro-parietal Junction (TPJ), N = 26. * is p < 0.05, ** is p < 0.005, *** is p < 0.0005 and n.s. indicates *not significant*. Error bars indicate standard error of mean.

This experiment replicated all the key findings from Experiment 1 (see Figures 2C-D, S6 and S7). Consistent with the domain-specific hypothesis of causal reasoning, the TPJ responded selectively to social causal content, whereas the PN responded selectively to physical causal content. Specifically, in both regions, responses were significantly greater for (i) causal than noncausal content within the preferred domain and (ii) causal content in the preferred than the nonpreferred domain, and iii) causal effects were greater in the preferred than nonpreferred domain (see Table 1 for stats).

As in Experiment 1, the Multiple Demand (MD) network showed a significant *reversed* effect of non-causal > causal only in the physical domain in the left hemisphere (p < 0.005; Figure S2), providing no evidence that this system is engaged on domain-general causal reasoning. We found similar results in the language network: most fROIs showed significant causal > non-causal effect in the social domain, and significant reversed effect of non-causal > causal in the physical domain (Figure S5). Thus, while the language network itself doesn’t show domain-general causal reasoning, linguistic stimuli can engage ToM network and the PN for domain-specific causal reasoning, aligning with the ‘Exportation Hypothesis’^34^. A whole-brain random effects group analysis for the conjunction of causal > non-causal contrasts in both physical and social domains did not reveal any domain-general neural substrate for causal reasoning (Figure 3, *bottom row*). A split-half GSS analysis on the conjunction of the two domain-specific contrasts revealed putative domain-general parcels in one half of the data, but none of them cross validated to show causal > non-causal responses across domains in the held-out data. Thus, across both fROI and whole-brain analyses, no brain region showed the response profile predicted by domain-general causal reasoning, replicating the pattern observed in Experiment 1.

**Figure 3.**
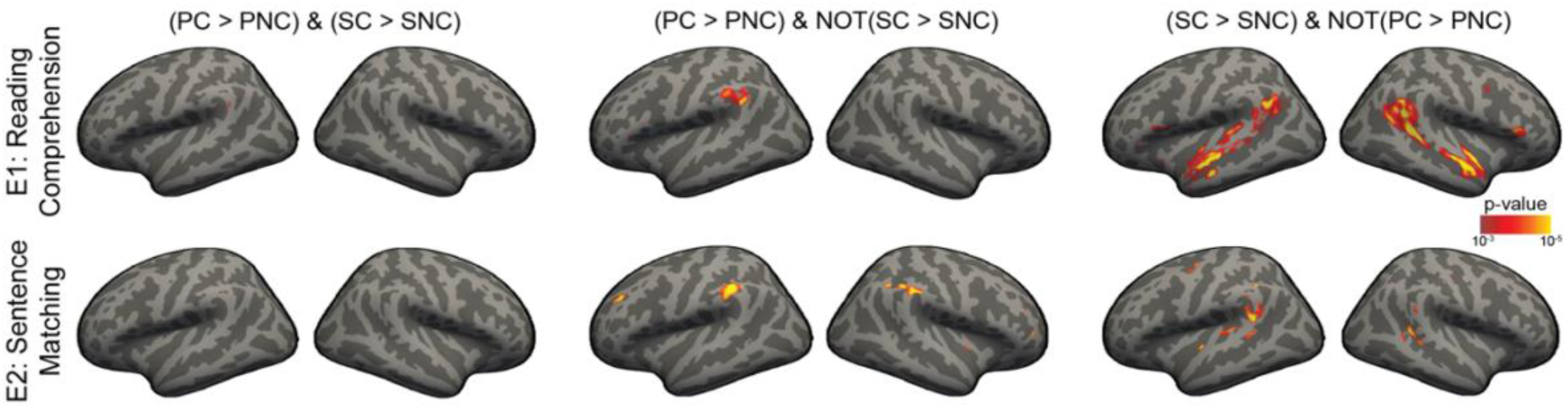
Random-effects analysis testing for domain-general causal reasoning. The top row shows significance maps for a whole-brain random effects (rfx) analysis in Experiment 1 (reading comprehension, N = 31, top) and Experiment 2 (sentence matching, N = 38, bottom) for the conjunction of causal > non-causal contrast in the physical and social domains (left); for causal > non-causal contrast in the physical domain, and *not* in the social domain (middle); for causal > non-causal contrast in the social domain, and *not* in the physical domain (right). All significance maps were thresholded at p < 0.001

Finally, although our study was not designed for a multivariate analysis, we nonetheless ran a brain-wide searchlight analysis on Experiments 1 and 2 (separately) to look for any brain regions showing stronger pattern correlations between the two causal conditions (i.e., r_(PC, SC)_) compared to the average correlations between the causal and non-causal conditions (i.e., average of r_(PC, PNC)_ and r_(SC, SNC)_), which would be suggestive of domain-general causal reasoning. We found no evidence for domain-generality in either experiment (Figure S8).

## Experiment 3: Generalization to causal preventative reasoning

Our findings from E1 and E2 consistently support the domain-specific hypothesis of causality, showing increased responses to causal over noncausal conditions only for the domain in question in domain-specific brain regions. Do these effects generalize to other forms of causal reasoning? To find out, we designed a third experiment (N = 17) with a new condition: causal reasoning about how an effect might have been *prevented,* as a test of the generality of our causality activations for different forms of causal reasoning. We included both physical causal preventative stimuli (e.g., in which *“The door hinge squeaked”* was paired with *“It should have been well oiled”*) and social causal preventative ones (e.g., in which *“She felt embarrassed to be there”* was paired with *“She shouldn’t have come without being invited”*).

This experiment again replicated all the key findings supporting domain-specific causal reasoning from Experiments 1 and 2 (see Figure 4): significantly higher responses in both the PN and TPJ for i) the causal condition in the preferred domain than for the noncausal condition in the preferred domain, ii) the causal condition in the preferred than nonpreferred domain, and iii) a significant interaction between physical/social x causal/noncausal (see Table 1). More importantly, we found higher responses to the causal preventative (CP) condition than to the non-causal condition in the preferred domain for both PN (p = 0.03) and TPJ (p < 0.0005), showing generalization to a different form of causal reasoning.

**Figure 4:**
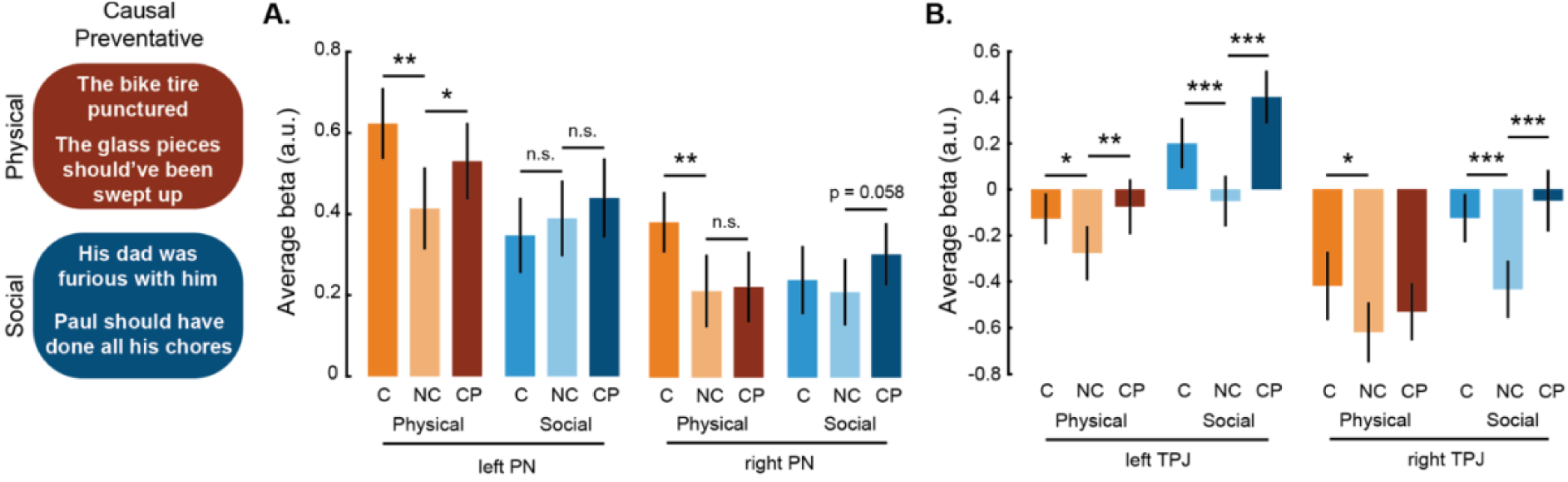
Experiment 3 (Causal Preventative Reasoning). This experiment constitutes an internal replication of the sentence matching task with causal and non-causal conditions in the physical and social domains, as well as a conceptual extension to causal preventative conditions (CP – in both physical and social domains; example stimuli on the left). (A) Bar plot showing the average activation to each of the six conditions in the left and right Physics Network (PN). (C) Similar bar plot in the left and right TPJ. * is p < 0.05, ** is p < 0.005, *** is p < 0.0005 and n.s. indicates *not significant*. Error bars indicate standard error of mean.

In both this experiment and in Experiment 2 (but not in Experiment 1) we observed a significantly higher response to the causal than non-causal condition for the non-preferred (physical) domain in the TPJ, albeit significantly smaller than for the preferred social domain. This raised at least two possibilities: that TPJ is engaged in causal reasoning across content domains, or that the non-preferred domain responses were driven in part by domain-general abstract relational reasoning not specific to causal reasoning. To test these possibilities, we also included stimuli testing reasoning about two abstract but non-causal relationships: part-whole and temporal order. We found that in the TPJ only the social (and not physical) causal condition was significantly higher than both the part-whole (p < 0.0005 and p < 0.0005 in lTPJ and rTPJ respectively) and temporal order (p < 0.0005 and p = 0.09 in lTPJ and rTPJ respectively) conditions (Figure S4). Thus, the TPJ is not engaged in domain-general causal reasoning more than other forms of abstract relational reasoning. We further found no evidence that that either the TPJ or PN play a broader role in domain-general abstract reasoning, as the response to part-whole and temporal order conditions were not significantly higher than the non-causal conditions (Figure S4). As in Experiments 1 and 2, the MD network showed no effect of causality in either domain (Figure S2) and no differential response to causal preventative versus non-causal conditions.

These results confirm and extend our previous findings on domain-specific causal reasoning in the human brain. Both PN and TPJ respond to the causal preventative condition in their respective domains, suggesting these regions support causal reasoning broadly.

## Discussion

Causal knowledge gives human beings the power not just to predict the future but to change it. In journals like this one, we ask “why” questions and offer “because” explanations across fields of inquiry. We isolate variables and assess the conditional probability of outcomes given interventions whether we are interested in how particles behave or how humans do. And in everyday life, we explain, intervene, and reason about counterfactual possibilities whether we are thinking about why a bicycle broke or why a child is upset. A single, unified system of causal reasoning seems to be at work. But our results suggest otherwise: The brain dedicates distinct, non-overlapping parts of the cortex to reasoning about events in the physical and social world.

Across three experiments, we find that reasoning about physical causes preferentially engages the Physics Network (previously shown to selectively process physical properties and events) whereas reasoning about social causes preferentially engages a region in the TPJ (previously shown to selectively process information about other people’s thoughts). Both cortical systems generalize beyond cause-and-effect inferences to causal-preventative reasoning, indicating their broad involvement in multiple forms of causal reasoning. Although several prior studies have addressed related questions^20,21,36–41^, none implemented a design that could test the domain specificity versus generality of causal reasoning in the human brain.

Other brain regions could in principle process causal information across social and physical domains, but our evidence does not support this hypothesis: First, the most famously domain-general cortical system, the multiple demand system, does not respond more strongly to causal than noncausal conditions. Second, brain-wide univariate analyses find no regions that respond more in causal than noncausal tasks equally across social and physical domains. Third, brain-wide multivariate analyses find no regions that produce patterns of response across voxels that distinguish causal versus noncausal conditions similarly for physical and social conditions. Taken together, our findings show that causal reasoning recruits domain-specific cortical regions and challenge the idea of a unified system specialized for causal reasoning across content domains.

The lack of evidence in our data for a domain-general mechanism for causal reasoning may reflect the limited spatial resolution of fMRI, which aggregates activity across hundreds of thousands of neurons within each voxel. Methods operating at finer spatial scales may in future reveal neural populations selectively engaged in causal reasoning across domains. Nevertheless, our findings constrain this possibility: if such populations exist, they are likely either smaller or less spatially segregated than those supporting many other cognitive functions^42^, including the domain-specific causal responses robustly identified across all three experiments.

The lack of a specialized region for causal reasoning is unexpected given how fundamental causal reasoning is to human cognition and given the prominence of domain-general accounts of causal reasoning in cognitive science^2,6^. Our findings are consistent, however, with developmental evidence showing that domain-specific causal representations emerge in infancy and that children develop distinct intuitive theories for physics and psychology. Our findings are also consistent with the established roles of the PN and TPJ. The TPJ is selectively engaged in theory of mind (ToM) reasoning^28,29^, of which a core component entails the causal inferences of attributing observed behavior to underlying goals, beliefs, and desires.

Causal reasoning offers a common framework for understanding how the PN and TPJ support cognition in the physical and social domains, respectively. Visual and auditory pathways provide representations of the agents, objects, and events in a scene, supplying the perceptual content from which richer interpretations can be constructed. The PN and TPJ organize this information into models of how objects and agents behave in the world. In the PN, forces and physical interactions provide the causal structure needed to explain object behavior; in the TPJ, goals, beliefs, and desires provide the causal structure needed to explain agent behavior^30,31^. These systems therefore allow us to move beyond registering what is present to understanding what is happening and why, enabling us to predict and intervene on future events in both the physical and social worlds.

Our study raises important questions for future research. First, many scenarios entail both physical and social phenomena, including cases in which a social effect has a physical cause (e.g. an earthquake causing people to fear for their lives), or a physical effect has a social cause (e.g. a multistory building collapsing because a corrupt public official decided not to enforce building codes). How is information integrated across domain-specific regions when multiple domains are implicated?^43^ Second, are fundamentally different computations entailed in reasoning about physical and social causes, or might similar algorithms be running in distinct brain regions that process different content domains? Perhaps most importantly, why does the fundamental ability to reason about causal events apparently not get its own cortical region, when so many other cognitive processes (including most recently logical reasoning)^44^ do?

The finding of distinct neural bases for physical and social causal reasoning is important not just for what it tells us about the functional organization of the brain, but for how it contributes to longstanding debates in philosophy and psychology. Many scholars have proposed a distinction between “reasons” and “causes”, arguing that human behavior is governed by the former not the latter^45,46^; others have claimed that insofar as reasons are efficacious, they are indistinguishable from causes^47^. Neural evidence may not have the last word in these philosophical disputes, but any attempt to understand human cognition must take such evidence into account.

## Methods

### fMRI data acquisition and pre-processing

All imaging was performed on a Siemens 3T Prisma scanner with a 32-channel head coil at the Athinoula A. Martinos Imaging Center at MIT. For each participant, a high-resolution T1-weighted anatomical image (MPRAGE: TR = 2.53 s; TE = 3.57 ms; a = 9°; FOV = 256 mm; Matrix = 256x256; Slice thickness = 1 mm; 176 slices; Acceleration factor = 2; 24 reference lines; BW=190Hz/pix) was collected in addition to whole-brain functional data using a T2*-weighted echo planar imaging pulse sequence (TR = 2 s; TE = 30 ms; a = 90°; FOV = 204 mm; Matrix = 102 x 102; Slice thickness = 2 mm; Voxel size = 2 x 2 mm in-plane; Slice gap = 0 mm; 66 slices).

Participants viewed stimuli through a mirror projected to a 12x16” screen behind the scanner, at a visual angle of approximately 14°x19°.

Preprocessing was done using FreeSurfer (https://freesurfer.net/). All other analyses were performed in MATLAB. fMRI data preprocessing included motion correction, slice time correction, linear fit to detrend the time series, and spatial smoothing with a Gaussian kernel (FWHM = 5 mm). Before smoothing the functional data, all functional runs were co-registered to the subject’s T1-weighted anatomical image. All analyses were performed in each subject’s native volume (except the searchlight analyses). The general linear model included the experimental conditions and 6 nuisance regressors based on the motion estimates (x, y, and z translation; roll, pitch, and yaw of rotation).

## Functional Localizers

Three fMRI “localizer” experiments were run to localize previously established functional Regions of Interest (fROIs) in each participant individually, thus accounting for variability in the exact location of each region across individuals.

### Physics (or Towers) localizer

Each participant performed two runs of an ‘intuitive physics’ fMRI localizer task previously used to functionally define the frontoparietal physics engine in the brain^23^. In this task, subjects viewed short movies (∼6 s) depicting unstable towers made of blue, yellow and white blocks created using Blender 2.70 (Blender Foundation). The tower was centered on a floor that was colored green on one half and red on the other half such that it would topple toward one of the halves if gravity were to take effect. Throughout the movie, the tower remained stationary while the camera panned 360° to reveal different views of the tower. Subjects viewed these movies and were instructed to report whether the block tower would fall on the red or green half of the floor (‘physics’ task), or whether there are more blue or yellow blocks in the tower (‘color’ task).

Each run of this localizer task consisted of 23 18 s blocks: 3 fixation-only blocks, 10 blocks each of the physics and the color task. Each 6 s movie was preceded by a text instruction displayed on the screen for 1 s which read either ‘where will it fall?’ (‘physics’ task) or ‘more blue or yellow?’ (‘color’ task) and was followed by a 2-s response period with a blank screen. This sequence was repeated twice within a block with the same task cue but different movies. The subjects responded by pressing one of two buttons on a response box for each alternative in a task. The mapping of the buttons to the response was switched for the second run to rule out the effects of specific motor responses on the observed neural activations. We used a physics task > color task contrast to functionally identify the frontoparietal physics regions in each subject individually.

### Theory of Mind (ToM) localizer

Participants were scanned on two runs of a standard theory of mind localizer (Jacoby et al, 2016) to identify brain regions previously implicated in inferences about others’ mental states^31^. Participants answered a true/false question after reading a short story describing a false representation that is either mentally held by a person (‘False Belief’ condition) or physically present on a map/photo (‘False Photo’ condition). Each run contained 5 False Belief and 5 False Photo stories. Each story was presented for 10s after which a true/false question about either the true state of the world or the false representation was presented for 4s, with a 12s inter-stimulus interval. The False Belief > False Photo contrast was used to identify ToM regions in each individual participant.

### Spatial Working Memory (SWM) localizer

Each participant completed two runs of a spatial working memory fMRI localizer task previously used to define the frontoparietal ‘multiple demand’ (MD) regions^33^. On each trial, subjects saw a series of 4x4 grids and kept track of either four or eight locations (‘easy’ and ‘hard’ trials, respectively) which had been marked on these grids either one or two at a time. At the end of each trial, subjects selected which of two grids was showing the cumulative locations indicated on the previous grids in a two-alternative forced-choice decision. Each run consisted of 48 eight-second trial blocks and four 16-second fixation blocks, with a palindromic ordering of hard and easy trials counterbalanced across runs. We used a hard > easy contrast to functionally identify MD regions in each subject individually.

### Language localizer

We used a task used to localize the language network in previous studies^48^. Briefly, we used a reading-based task that contrasted English sentences and lists containing unconnected, pronounceable non-words in a standard block design. Each block contained 12-word sequences presented one word at a time followed by a prompt for the subject to push a button (to make sure they are paying attention). The participants were instructed to read each word or non-word and press a button when the image of a hand pressing a button was displayed. Each block lasted for 6s: 100 ms of blank screen, then each of the 12 words/non-words displayed sequentially for 450 ms, then the button press image for 400 ms, followed by a final 100 ms of blank screen. Each participant performed 2 runs of this localizer task where each run contained 16 such blocks (8 English sentences and 8 non-word lists). The block orders were counterbalanced across runs.

## Experiment 1: Reading Comprehension Task

The experiment design and analyses were pre-registered here: https://osf.io/d7s29 (and consolidated in https://osf.io/j2qt8). As in Experiment 1, the domain-specific causality hypotheses in PN and ToM regions and the univariate analyses therein follows from the pre-registration, whereas the univariate analysis within MD (to test domain general causality hypothesis) and grou-level RFX and surface searchlight MVPA are exploratory.

### Participants

Were recruited from Massachusetts Institute of Technology and the surrounding Greater Boston area (N = 31, 15 females; aged 19-58 with a median age of 24). All participants had normal or corrected-to-normal vision and gave informed consent to the experimental protocol approved by the Massachusetts Institute of Technology (MIT) Committee on the Use of Humans as Experimental Subjects (COUHES #0403000096).

### Experiment design

In this experiment, subjects read short stories, each describing an event that entailed a social cause or physical cause (SC, PC), or that described socially or physically noncausal (spatial or constitutive) relations (SNC, PNC).

In each block of this task, a short story corresponding to one of the stimulus conditions was displayed for 13 seconds. Following the short story, a multiple-choice question was presented for 9 seconds below the short story. Thus, each stimulus block lasted 22 seconds. The participants were instructed to choose the correct answer by pressing the appropriate button on a button box. The complete stimulus set contained 16 stories for each of the four conditions. All four conditions were shown four times in each run and organized across the four runs such that they appear in every serial position an equal number of times. Each run consisted of 16 stimulus blocks and five 14s fixation blocks uniformly interleaved within the run, and short fixation blocks between stories lasting for 2s acting as transition periods between stimulus blocks. The fixation blocks showed a white cross in the center of a gray screen.

All verbal stimuli were matched across conditions for grade level, word count, and character count using online tools (https://readabilityformulas.com/readability-scoring-system.php; https://datayze.com/readability-analyzer). Additionally, stimuli were behaviorally tested prior to their use in fMRI experiments to approximately match conditions for difficulty (task accuracy and reaction times). Task performance inside the scanner (Figure S1) also confirmed our stimulus norming procedure, whereby both task accuracy and reaction times did not show any consistent trends across domains.

## Experiment 2: Sentence Matching Task

The experiment design and analyses were pre-registered here: https://osf.io/s54tb (and consolidated in https://osf.io/j2qt8). The domain-specific causality hypotheses in PN and ToM regions and the univariate analyses therein follows from the pre-registration, whereas the univariate analysis within MD (to test domain general causality hypothesis) and group-level RFX and surface searchlight MVPA are exploratory.

### Participants

Were recruited from Massachusetts Institute of Technology and the surrounding Greater Boston area (N = 38, 21 females; aged 20-58 with a median age of 24.5). All participants had normal or corrected-to-normal vision and gave informed consent to the experimental protocol approved by the Massachusetts Institute of Technology (MIT) Committee on the Use of Humans as Experimental Subjects (COUHES #0403000096). We collected data from more participants than our pre-registration indicates (N = 18), but our results are qualitatively similar with this smaller participant pool.

### Experiment design

This experiment is a within-subjects, 2 by 2 design with two orthogonal factors: Causality (causal versus descriptive) and Domain (physical versus social). We tested our three hypotheses by scanning participants with fMRI where they matched causes to their effects or matched descriptions of the same entity.

Each block of the experiment consisted of an array of 8 sentences, from one of the four conditions: physical causal (PC), social causal (SC), physical non-causal/descriptive (PNC), or social non-causal/descriptive (SNC). Specifically, each block showed four images of ice cream scoops (top) and four images of ice cream cones (bottom) that participants had to match, progressing from left to right (indicated by a red line, see Figure 2B). To induce the subjects to think about the causal or descriptive nature of our verbal stimuli, they were asked to match the phrases written in the ice cream scoops to the phrases written in the ice cream cones. Prior to seeing each block of ice cream stimuli, participants read the instructions (for four seconds) to “match the causes to their effects” or “match the descriptions of the same entity.” The participant chose the scoop that corresponded to the cone that is highlighted by the red bar underneath by pressing the corresponding button (buttons 1-4).

Each block lasted 20 seconds, but the participants were allowed to match the items at their own pace within this duration. The complete stimulus set contained 64 pairs of sentences (arranged in 16 four-pair trials) for each of the four conditions. Within each block the scoop images were shuffled randomly. Each run consisted of 8 stimulus blocks and three fixation blocks – one at the beginning, middle, and end of each run. Each of the four conditions was shown twice in each run, with conditions ordered palindromically within each run. Across runs there was an equal number of trials of each condition in each serial position within a run. The fixation blocks lasted 18 seconds and displayed a white fixation cross in the center of a gray screen.

The stimuli were matched across conditions for grade level, word count, and character count using an online tool (https://datayze.com/readability-analyzer). Additionally, online surveys were created using Qualtrics and distributed via Prolific prior to their use in fMRI to approximately match conditions for difficulty in terms of task accuracy and response time. Task performance inside the scanner (Figure S1) also confirmed our stimulus norming procedure, whereby both task accuracy and reaction times did not show any consistent trends across domains.

## Experiment 3: Sentence Matching Task with additional conditions

The experiment design and analyses were pre-registered here: https://osf.io/d7s29 (and consolidated in https://osf.io/j2qt8). As in Experiment 1, the domain-specific causality hypotheses in PN and ToM regions and the univariate analyses therein follows from the pre-registration, whereas the univariate analysis within MD (to test domain general causality hypothesis) is exploratory.

### Participants

Were recruited from Massachusetts Institute of Technology and the surrounding Greater Boston area (N = 17, 10 females; aged 19-58 with a median age of 23). All participants had normal or corrected-to-normal vision and gave informed consent to the experimental protocol approved by the Massachusetts Institute of Technology (MIT) Committee on the Use of Humans as Experimental Subjects (COUHES #0403000096).

### Experimental design

The design is similar to the Sentence Matching task (E2) described above but with 4 additional conditions for a total of 8 conditions. Specifically, in addition to the physical and social causal and non-causal conditions from before, we included physical and social causal preventative (CP) conditions to generalize our results to a new kind of causal reasoning, and two abstract relational reasoning conditions – part/whole (PW) and temporal order (TO) – to test whether stronger responses to the causal condition in the non-preferred domain (for example, physical causal > non-causal in TPJ) is actually driven by causal reasoning or any other kind of abstract relational reasoning.

Each block in this experiment consisted of stimuli from one of the eight conditions. As in E1, each block showed four images of ice cream scoops (top) and four images of ice cream cones (bottom) that participants were to match, progressing from left to right (indicated by a red line, see Figure 2B). To induce subjects to think about the causal, descriptive, and other abstract relationships of our stimuli, they were asked to match the phrases written in the ice cream scoops to the phrases written in the ice cream cones. Prior to seeing each block the stimuli, participants read the instructions (for four seconds) to “Match the EFFECTS (in cones) to the CAUSES (in scoops)”, “Match the EFFECTS (in cones) that could have been prevented by the interventions (in scoops)”, “Match the sentences (in cones and scoops) that describe the same entity.”, “Match the events (in cones) to the events that precede them (in scoops)”, or “Match the parts (in cones) to their respective wholes (in scoops).” The participant chose the scoop that corresponded to the cone highlighted by the red underline by pressing a button.

Each block lasted for 20 seconds (not including the preceding 4 second instruction block), but participants could match the items at their own pace within that window. The complete stimulus set contained 32 pairs of sentences for each of the eight conditions. Each run consisted of 12 stimulus blocks (8 conditions all matched for difficulty, grade level and word count + 4 conditions from E1 presented as is for internal replication) and three fixation blocks (one at the beginning, middle, and end of each run). The fixation blocks lasted 20 sec and showed a white cross in the center of a gray screen.

## Definition of functional Regions of Interest (fROIs)

### General Method

Functional regions of interest (fROIs) were defined separately within each participant as the intersection of voxels reaching significance in the relevant contrast and the relevant anatomical constraint parcel obtained from prior publications (PN parcels^25^, ToM parcels from https://saxelab.mit.edu/use-our-theory-mind-group-maps/; and MD and language network parcels from https://www.evlab.mit.edu/resources). We transformed group-level parcels into the subject’s native space, and then selected voxels falling within these parcels based on their contrast significance in the localizer runs. Specifically, we selected voxels that showed significant contrast at p < 0.001 and if there were fewer than 50 voxels with this threshold, we reduced the threshold to p < 0.05. Subsequent analyses were performed in these selected voxels using functional data from the experimental runs (E1, E2 and E3). Table S1 shows the distribution of experiments and localizer tasks across participants.

## Analyses methods

### fROI analysis (univariate)

The univariate analyses in the PN, TPJ, MD, and the Language Network were conducted on the fMRI data from the main experiments, which was independent of the data used to localize these fROIs. In each experiment and for each participant, we computed the average activations across all functionally localized voxels within each ROI for each experimental condition. To compare activations for causal and non-causal conditions within a domain, we performed a one-tailed Wilcoxon signed-rank test (using MATLAB *signrank* function). Further, we characterized interaction between domain and causality within each fROI using repeated measures, mixed-effects ANOVA (MATLAB *anovan* function) with participants as random factors, and domain and causality as fixed factors.

### Random effects analysis

Individual subject domain-specific causal > non-casual contrast maps were transformed from subject-native space to *fsaverage* using FreeSurfer’s mri_vol2vol function with affine registration matrices computed during preprocessing. Specifically, contrast effect size (ces) and variance (cesvar) maps were transformed for all subjects in both sentence matching and reading comprehension tasks. In each experiment, the transformed contrast maps were concatenated across subjects, and a one-sample group-level random-effects (RFX) general linear model was performed using FreeSurfer’s mri_glmfit with the --osgm flag. This approach treats subjects as a random effect and tests whether mean activation across the group significantly differs from zero, providing conservative group-level inference (Figure S3).

To identify domain-general regions (i.e, regions showing significant activation for both physical and social causality), we implemented a logical AND conjunction analysis. Specifically, at each voxel, we extracted the minimum significance value between the two domain-specific RFX significance maps, such that only voxels significant in both contrasts were retained in the overlap map. To control for multiple comparisons and remove spurious isolated voxels, we applied cluster-level correction by thresholding the overlap map at -log₁₀(p) = 3.0 (p < 0.001, uncorrected) and retaining only clusters comprising a minimum of 50 contiguous voxels (on a volume-based 26-connectivity analysis).

### Multi-voxel pattern analysis

To examine how representational structure varies across cortex, we performed a surface-based searchlight analysis within each experiment. For each subject and at each cortical vertex, we computed pairwise correlations between all four task conditions (PC, PNC, SC, SNC) within a neighborhood centered at that vertex. Neighborhoods were defined using 10mm geodesic distance on the individual subject’s surface, ensuring spatially contiguous voxels while respecting cortical anatomy. At each searchlight location, we extracted a 4×4 correlation matrix from the condition-wise beta estimates within the neighborhood, then stored the six unique pairwise correlations: PC-PNC, PC-SC, PC-SNC, PNC-SC, PNC-SNC, and SC-SNC. This process was repeated for all vertices across the hemispherical surface, yielding vertex-wise maps of representational structure.

To characterize representational structure, we computed correlation differences that indexed three organizational principles. Domain-generality was quantified as the correlation between physical and social causal conditions (PC-SC) minus the average cross-causality correlations within each domain (PC-PNC and SC-SNC), reflecting whether the causal conditions are more similar across domains than they are to the non-causal conditions within their respective domains. A lack of domain-generality in this analysis could also imply that the analysis is not powerful enough to reveal an effect if it was indeed present. So as controls, we also tested two additional organizational principles using the same three pair-wise correlations: i) correlation between physical conditions (PC-PNC) minus the average of correlations (PC-SC) and (SC-SNC), and ii) correlation between social conditions (SC-SNC) minus the average of correlations (PC-SC) and (PC-PNC). We did not have strong predictions for these organizational principles except that our univariate results pointed to ToM and PN respectively as potential substrates. The pairwise correlations were Fisher Z-transformed before computing the difference and each subject’s correlation difference maps were transformed to standard space (fsaverage).

At the group level, we performed one-sample t-tests on the correlation differences across subjects. To account for multiple comparisons, we employed a permutation-based cluster correction procedure (1,000 permutations) using sign-flipping of individual subject maps. For each permutation, we recomputed group-level t-statistics and identified clusters of contiguous vertices using graph-based connectivity. The cluster-level threshold was defined as the 95th percentile of the maximum cluster size distribution across permutations. Only clusters exceeding this threshold in the real data were retained for interpretation, ensuring family-wise error control at p < 0.05. The resultant maps are shown in Figure S8.

### Surface visualization

Statistical maps were projected onto the fsaverage cortical surface for both hemispheres using FreeSurfer’s *mri_vol2surf* function with nearest-neighbor interpolation and a projection fraction range of -0.1 to 1.0 (normalized units from white matter to pial surface) to reduce spatial noise artifacts. Surface maps were visualized using FreeView (Figures 3, S3 and S8).

## Data availability statement

The fMRI data (anonymized) generated during the study will be deposited in a public repository upon publication.

## Acknowledgements

We thank Rebecca Saxe, Evelina Fedorenko, the members of the Kanwisher Lab and Early Childhood Cognition Lab at MIT for their valuable feedback during the study. We thank Atsushi Takahashi and Steven Shannon of the Athinoula A. Martinos Imaging Center at the McGovern Institute for Brain Research, MIT for their fMRI expertise. Jessica Chomik-Morales would like to acknowledge the Post-Baccalaureate Research Scholars Program, Halis Family Foundation, and Gene Stark for their support.

## Author Information

### Author Contributions

R.T.P., J.C.-M., L.S., and N.K. conceived and designed the study; R.T.P. and J.C.-M. collected data; R.T.P. and J.C.-M. performed the univariate analyses; R.T.P. performed the random effects and multivariate analyses; L.S. and N.K. acquired funding and supervised the study; R.T.P. wrote the original draft of the manuscript with N.K.’s inputs. All authors contributed to manuscript preparation.

## Extended data figures and tables

**Figure S1:**
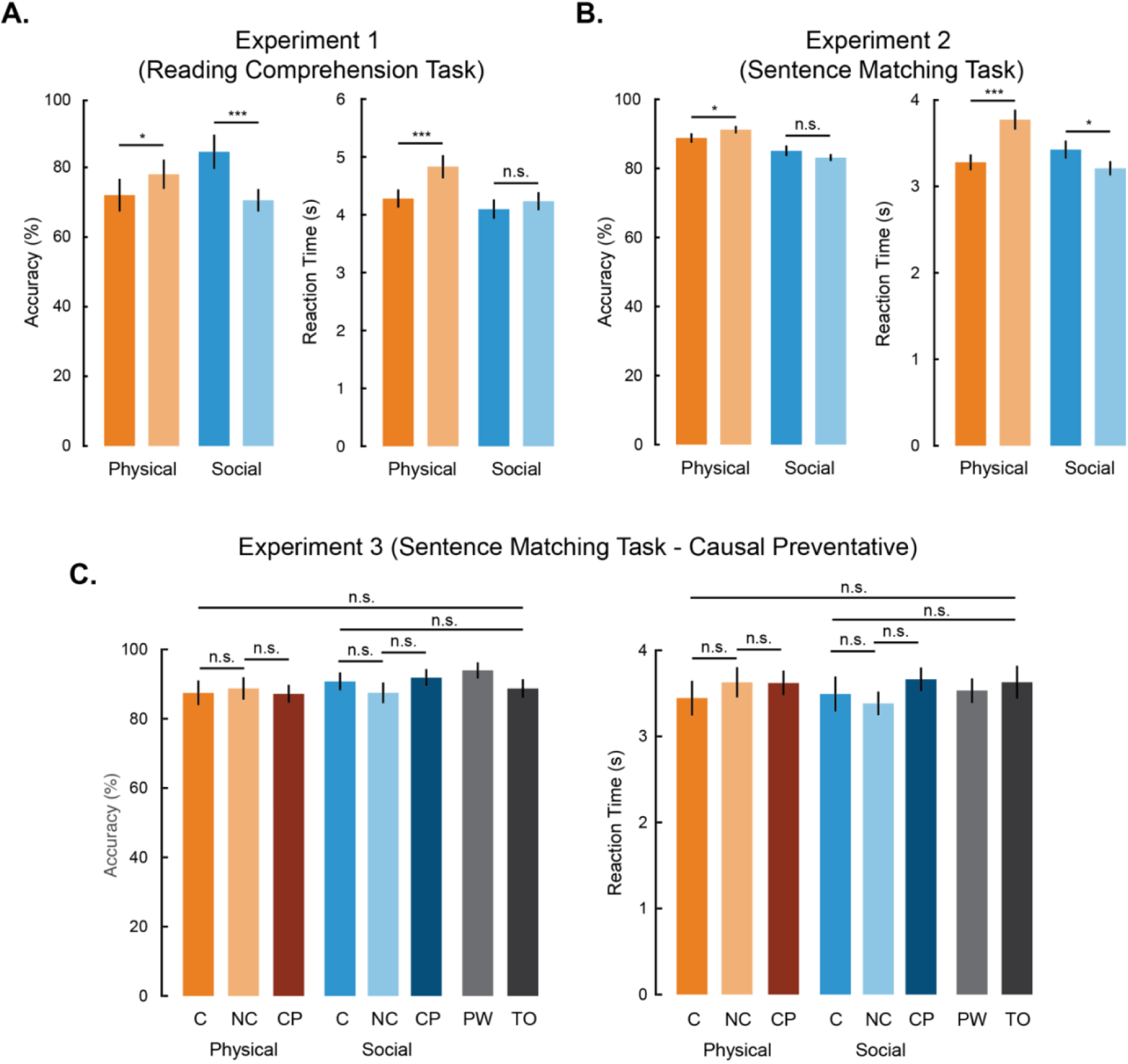
Behavioral performance of participants in the three experiments. (A) Average accuracy (% correct; left) and average reaction time (in seconds; right) across participants for all the four conditions in Experiment 1. (B) and (C) similar plots for Experiments 2 and 3. * is p < 0.05, ** is p < 0.005, *** is p < 0.0005 and n.s. is not significant on a wilcoxon ranksum test. Error bars are standard error of mean.

**Figure S2:**
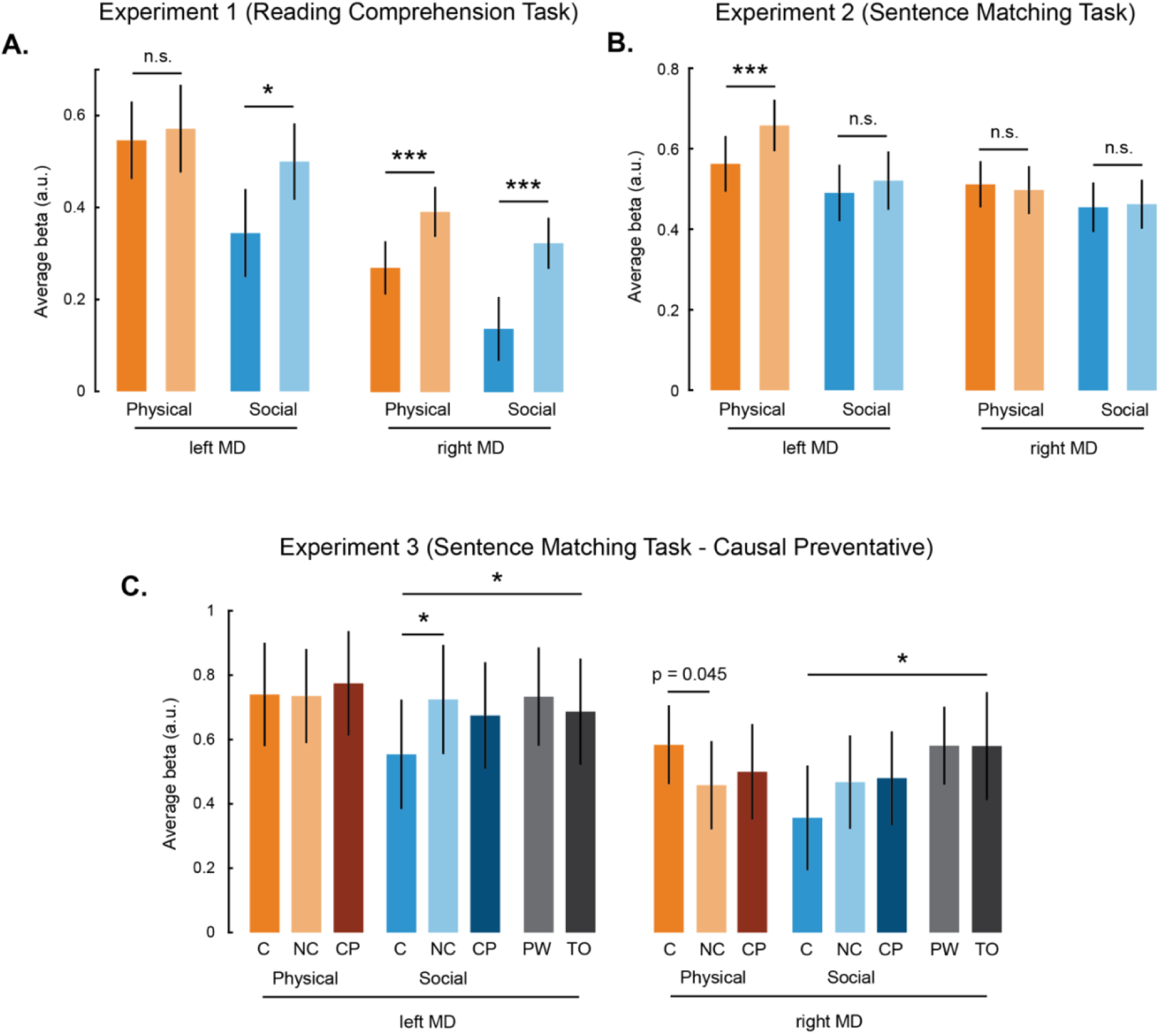
Responses the Multiple Demand (MD) network. (A) Bar plot showing the average fMRI response to each of the four conditions in the left and right hemispheres of the MD network in Experiment 1. Darker colors are causal and lighter colors are non-causal conditions. (B) is a similar bar plot for Experiment 2. Darker colors are causal and lighter colors are non-causal conditions. (C) is for Experiment 3 showing average fMRI responses to Causal (C), Non-causal (NC), Causal Preventative (CP), Part-Whole (PW), and Temporal Order (TO) conditions. * is p < 0.05, ** is p < 0.005, *** is p < 0.0005 and n.s. is not significant for a one-sided Wilcoxon ranksum test. Error bars are standard error of mean.

**Figure S3:**
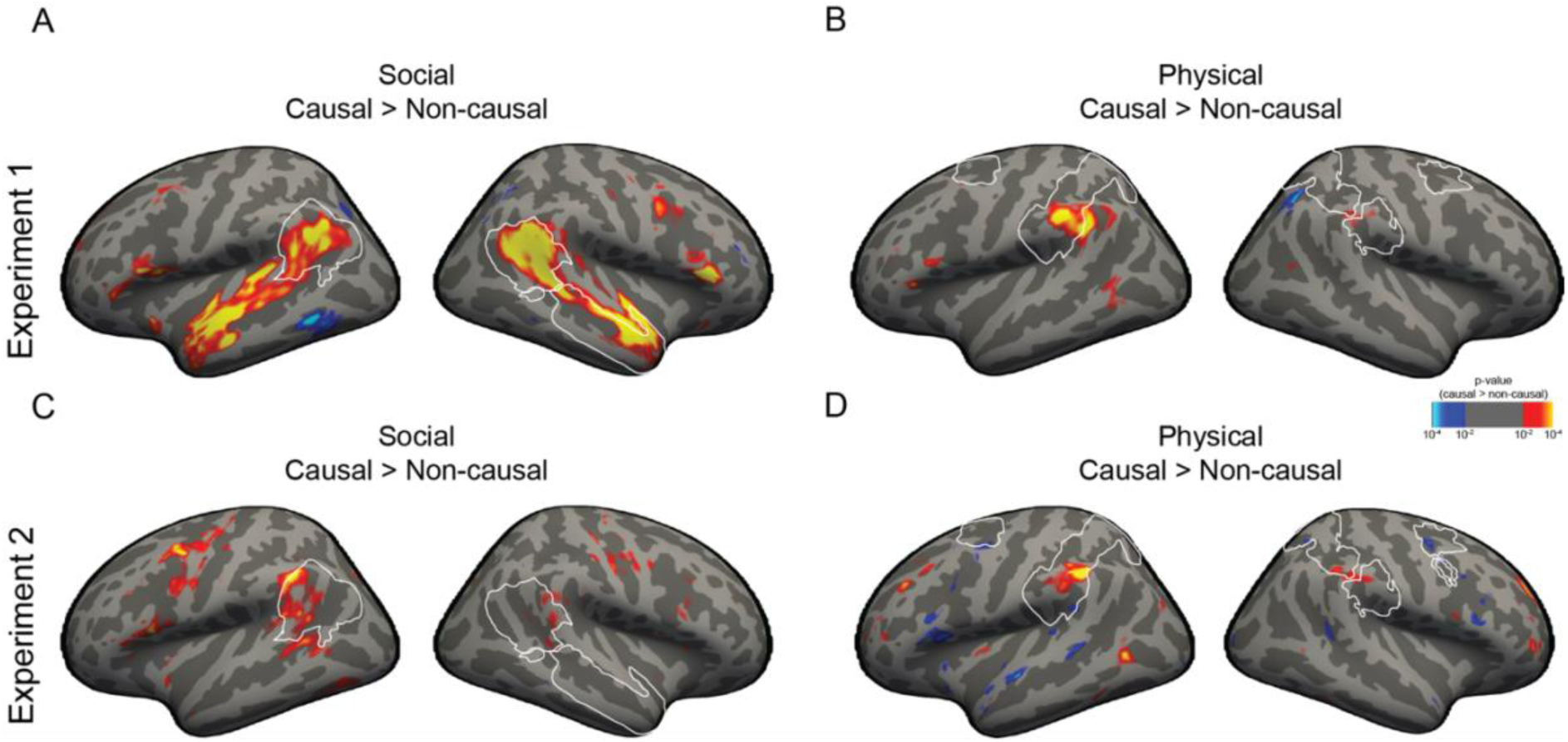
Group-level random effects (rfx) analysis. (A) and (B) are significance maps in Experiment 1 for social and physical causality respectively shown on inflated cortical surface. (C) and (D) similar domain-specific significance maps for Experiment 2. The white outlines indicate bilateral TPJ and right hemisphere STS anatomical masks on the social causality RFX maps, and bilateral PN anatomical masks on the physical causality RFX maps.

**Figure S4:**
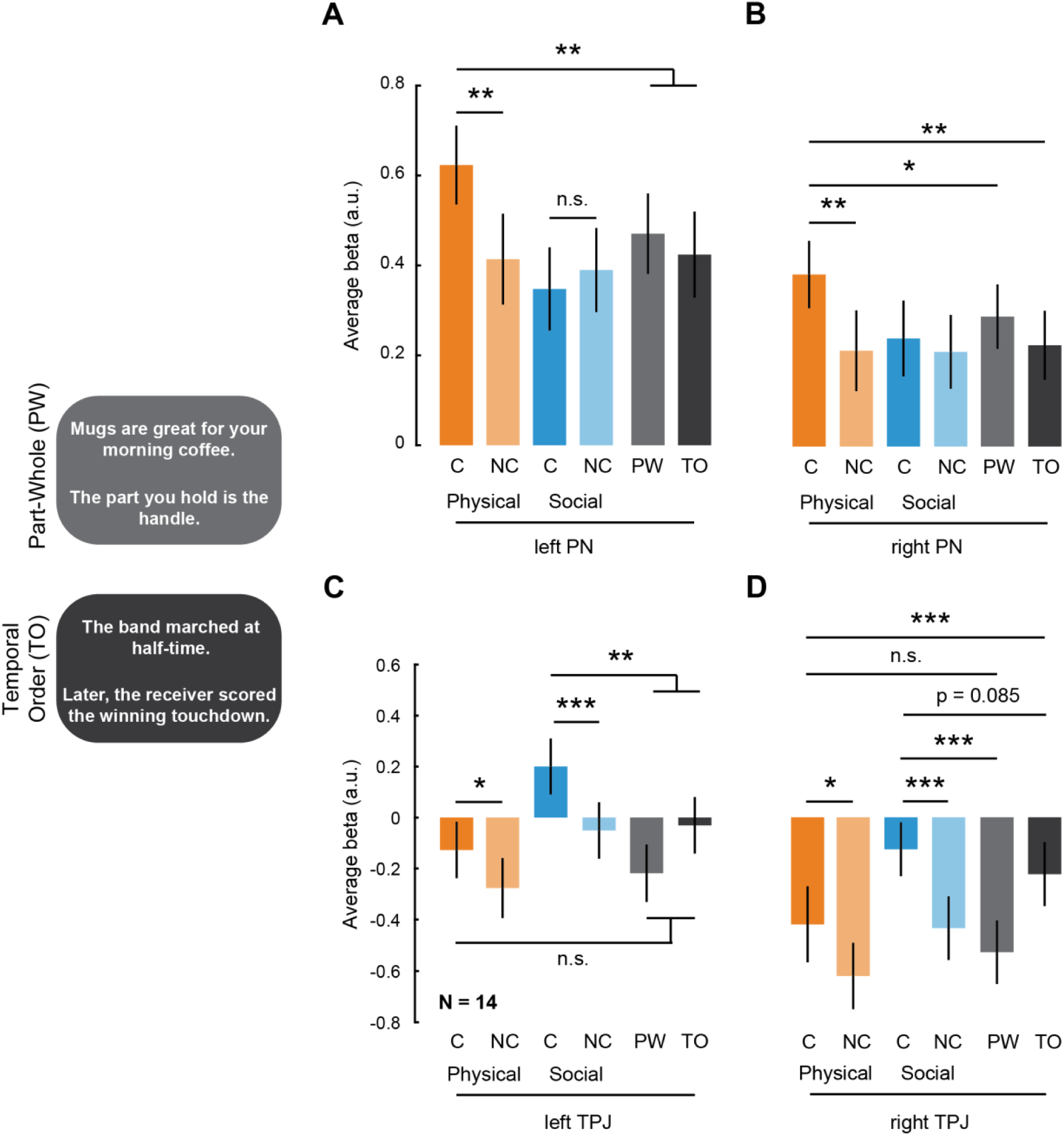
Responses to abstract relational reasoning in Experiment 3. Example Part-Whole (PW) and Temporal Order (TO) reasoning stimuli. (A) and (B) are bar plots showing average activation for causal and non-causal conditions in physical and social domains, and part-whole and temporal order conditions in left and right PN respectively. (C) and (D) are similar bar plots for left and right TPJ respectively. * is p < 0.05, ** is p < 0.005, *** is p < 0.0005 and n.s. is not significant for a one-sided Wilcoxon ranksum test. Error bars are standard error of mean.

**Figure S5:**
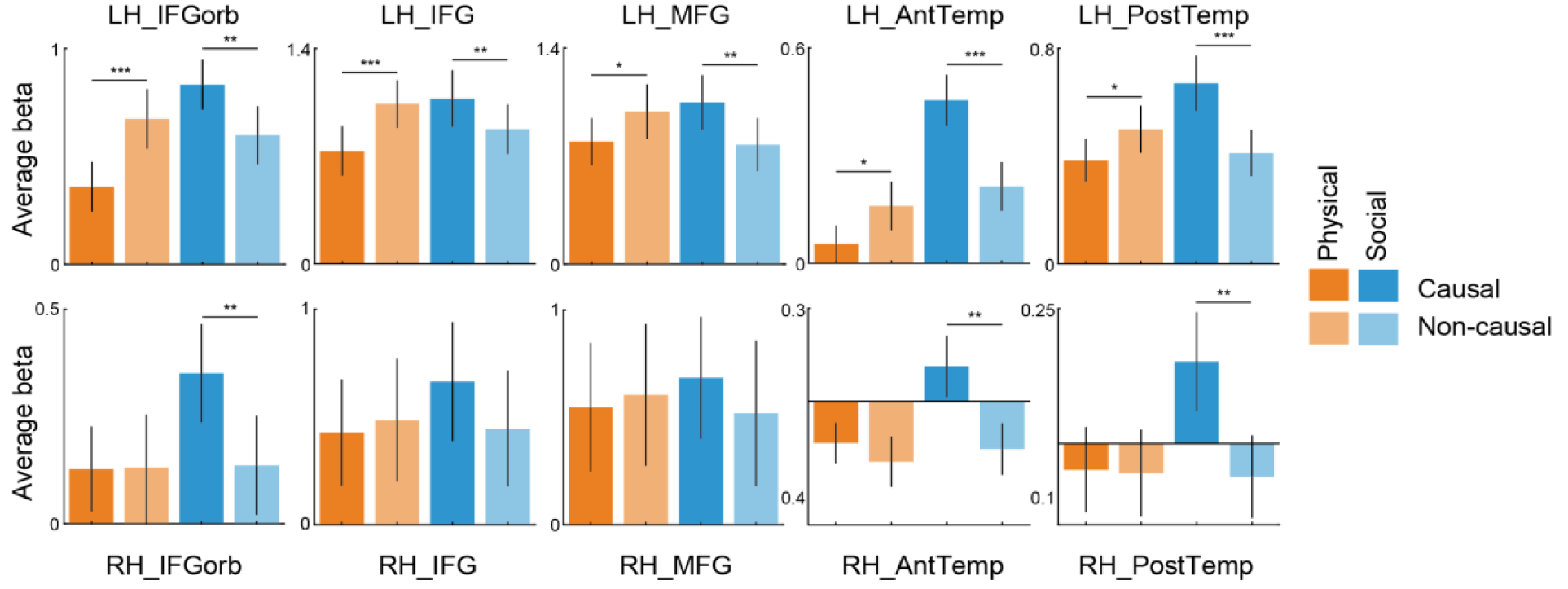
Language fROI responses in Experiment 2. We used a standard language localizer contrasting sentences and non-words to localize the language network in individual participants (N = 18). The bar plots depict average response across all participants for each of the four experimental conditions within each language region. * is p < 0.05, ** is p < 0.005, and *** is p < 0.0005 for a one-sided Wilcoxon ranksum test. Error bars are standard error of mean.

**Figure S6:**
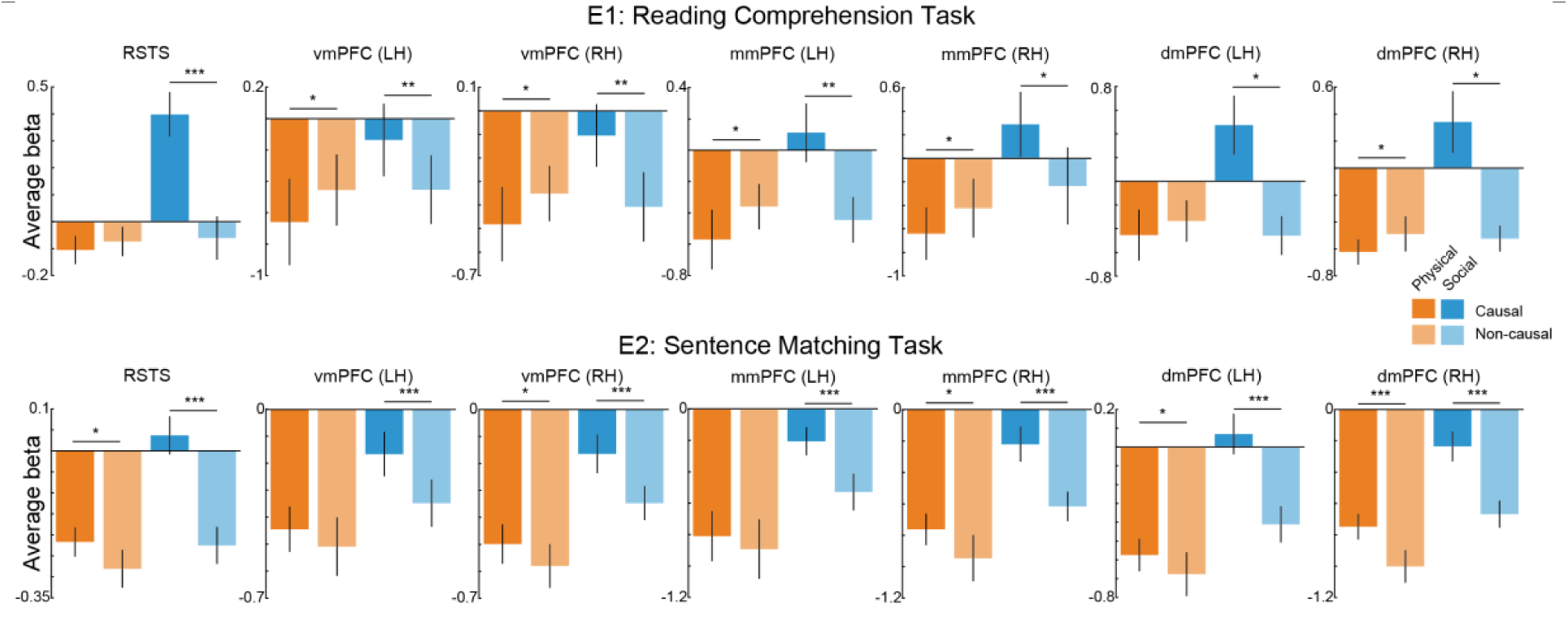
Responses in different ROIs of the Theory of Mind system in Experiments 1 (top row) and 2 (bottom row). The bar plots depict average response across all participants for each of the four experimental conditions within each ToM region localized using an independent localizer task. * is p < 0.05, ** is p < 0.005, and *** is p < 0.0005 for a one-sided Wilcoxon ranksum test. Error bars are standard error of mean.

**Figure S7:**
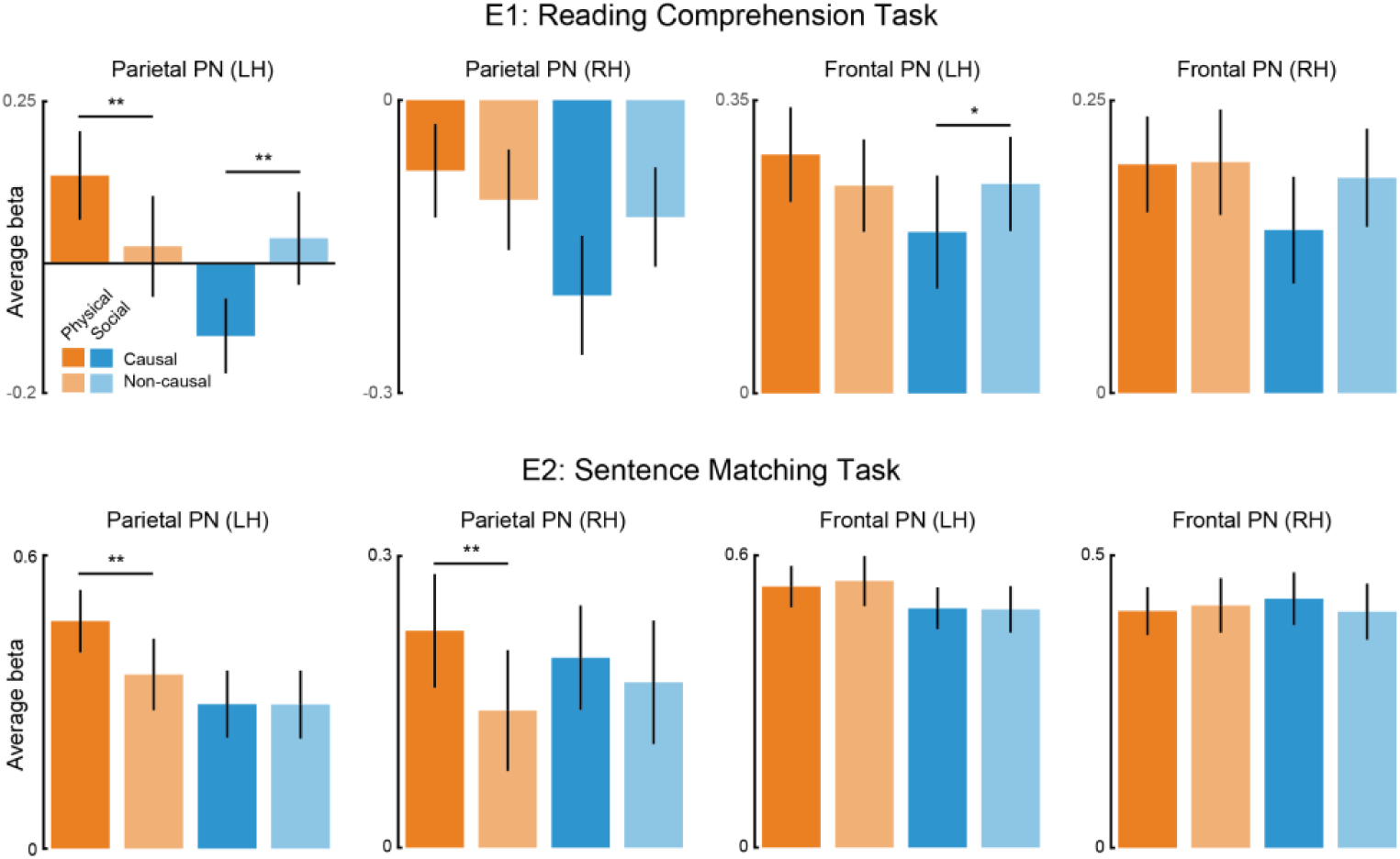
Responses in sub-regions of the PN in Experiments 1 (top row) and 2 (bottom row). The bar plots show the same data in Figures 1 and 2 for the PN except now they are split into different sub-regions of the bilateral PN. * is p < 0.05 and ** is p < 0.005 for a one-sided Wilcoxon ranksum test. Error bars are standard error of mean.

**Figure S8:**
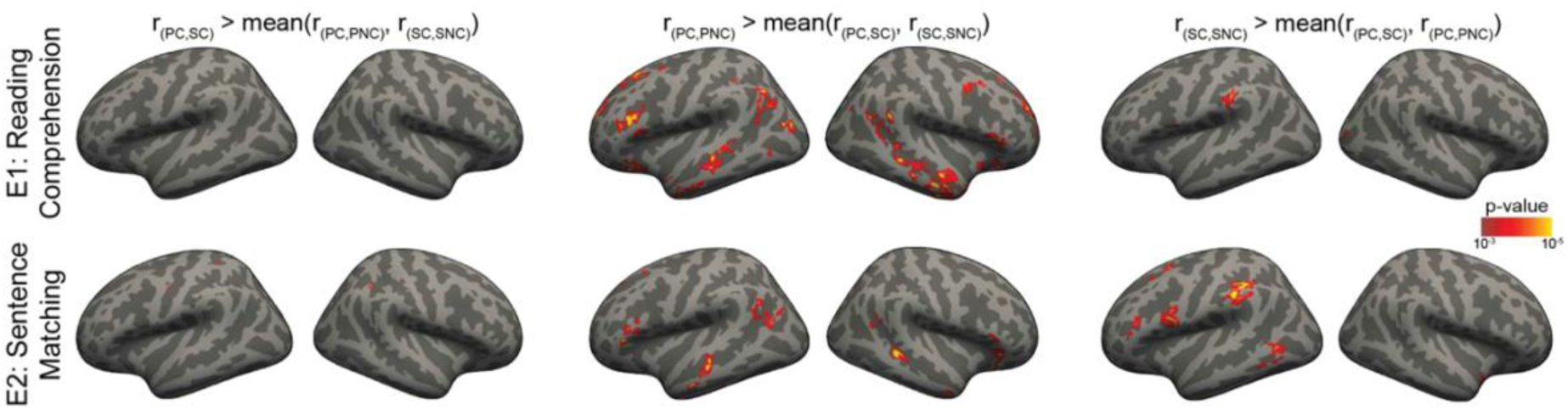
Multi-Voxel Pattern Analysis (MVPA) significance maps for in Experiments 1 (top row) and 2 (bottom row). We quantified domain-generality (left column) as the correlation between physical and social causal conditions (PC-SC) minus the average cross-causality correlations within each domain (PC-PNC and SC-SNC), reflecting whether the causal conditions are more similar across domains than they are to the non-causal conditions within their respective domains. To test whether we had sufficient statistical power in our test of domain generality, we also tested two additional organizational principles using the same three pair-wise correlations (for comparable statistical power): i) correlation between physical conditions (PC-PNC) minus the average of correlations (PC-SC) and (SC-SNC) in the middle column, and ii) correlation between social conditions (SC-SNC) minus the average of correlations (PC-SC) and (PC-PNC) in the right column. We did not have strong predictions for this control analysis, but we note that the results can be interpreted to show that brain regions that “do not know the difference” between physical causal and physical non-causal conditions are the social regions and vice versa.

**Table S1:** Tasks and Localizers run in each participant.

| **Subject ID** | **Reading**<br>**Comprehen**<br>**sion Task**<br>**(E1)** | **Sentence**<br>**Matching**<br>**Task (E2)** | **Sentence**<br>**Matching**<br>**Task (E3)** | **Theory of**<br>**Mind**<br>**Localizer** | **Spatial**<br>**Working**<br>**Memory** | **Tower**<br>**localizer** |
| --- | --- | --- | --- | --- | --- | --- |
| kancause03 | 1 |  |  |  |  | 1 |
| kancause04 | 1 |  |  |  |  | 1 |
| kancause05 | 1 |  |  |  |  | 1 |
| kancause06 | 1 |  |  |  |  | 1 |
| kancause07 | 1 |  |  |  |  | 1 |
| kancause08 | 1 |  |  |  |  | 1 |
| kancause09 | 1 |  |  |  |  | 1 |
| kancause10 | 1 |  |  |  |  | 1 |
| kanIC04 |  | 1 |  | 1 |  | 1 |
| kanIC05 |  | 1 |  | 1 |  | 1 |
| kanIC06 |  | 1 |  | 1 |  | 1 |
| kanIC07 |  | 1 |  | 1 |  | 1 |
| kanIC08 |  | 1 |  | 1 |  | 1 |
| kanIC09 |  | 1 |  | 1 |  | 1 |
| kanIC10 |  | 1 |  | 1 | 1 | 1 |
| kanIC11 |  | 1 |  | 1 | 1 | 1 |
| kanIC12 |  | 1 |  | 1 | 1 | 1 |
| kanIC13 |  | 1 |  | 1 | 1 | 1 |
| kanIC14 |  | 1 |  | 1 | 1 | 1 |
| kanIC15 |  | 1 |  | 1 | 1 | 1 |
| kanIC16 |  | 1 |  | 1 | 1 | 1 |
| kanIC17 |  | 1 |  | 1 | 1 | 1 |
| kanIC18 |  | 1 |  | 1 | 1 | 1 |
| kanIC19 |  | 1 |  | 1 | 1 | 1 |
| kanIC20 |  | 1 |  | 1 | 1 | 1 |
| kanIC21 |  | 1 |  | 1 | 1 | 1 |
| kanABIC03 | 1 | 1 | 1 | 1 | 1 | 1 |
| kanABIC04 | 1 |  | 1 |  | 1 | 1 |
| kanABIC05 | 1 | 1 | 1 | 1 | 1 | 1 |
| kanABIC06 | 1 |  | 1 |  | 1 | 1 |
| kanABIC07 | 1 | 1 | 1 | 1 | 1 | 1 |
| kanABIC08 | 1 | 1 | 1 | 1 | 1 | 1 |
| kanABIC09 | 1 |  | 1 |  | 1 | 1 |
| kanABIC10 | 1 | 1 |  |  | 1 | 1 |
| kanABIC11 | 1 | 1 |  |  |  | 1 |
| kanABIC12 | 1 | 1 |  |  | 1 | 1 |
| kanABIC13 | 1 | 1 |  |  | 1 | 1 |
| kanABIC14 | 1 | 1 |  |  | 1 | 1 |
| kanABIC15 | 1 | 1 | 1 | 1 |  | 1 |
| kanABIC16 | 1 | 1 |  |  | 1 | 1 |
| kanABIC17 | 1 | 1 |  |  | 1 | 1 |
| kanABIC18 | 1 | 1 |  |  | 1 | 1 |
| kanABIC19 | 1 | 1 | 1 | 1 | 1 | 1 |
| kanABIC20 | 1 | 1 | 1 | 1 | 1 | 1 |
| kanABIC21 | 1 | 1 |  |  | 1 | 1 |
| kanABIC22 | 1 | 1 |  |  | 1 | 1 |
| kanABIC23 | 1 | 1 |  |  |  | 1 |
| kanABIC24 | 1 | 1 |  |  | 1 | 1 |
| kanABIC25 | 1 | 1 | 1 | 1 | 1 | 1 |
| kanABIC26 |  |  | 1 | 1 |  | 1 |
| kanABIC27 |  |  | 1 | 1 |  | 1 |
| kanABIC28 |  |  | 1 | 1 |  | 1 |
| kanABIC29 |  |  | 1 | 1 |  | 1 |
| kanABIC30 |  |  | 1 | 1 |  | 1 |
| kanABIC31 |  |  | 1 | 1 |  | 1 |

